# Evaluating cancer cell line and patient-derived xenograft recapitulation of tumor and non-diseased tissue gene expression profiles *in silico*

**DOI:** 10.1101/2023.04.11.536431

**Authors:** Avery S. Williams, Elizabeth J. Wilk, Jennifer L. Fisher, Brittany N. Lasseigne

**Affiliations:** The Department of Cell, Developmental and Integrative Biology, Heersink School of Medicine, The University of Alabama at Birmingham, Birmingham, Alabama, USA

## Abstract

Preclinical models like cancer cell lines and patient-derived xenografts (PDXs) are vital for studying disease mechanisms and evaluating treatment options. It is essential that they accurately recapitulate the disease state of interest to generate results that will translate in the clinic. Prior studies have demonstrated that preclinical models do not recapitulate all biological aspects of human tissues, particularly with respect to the tissue of origin gene expression signatures. Therefore, it is critical to assess how well preclinical model gene expression profiles correlate with human cancer tissues to inform preclinical model selection and data analysis decisions. Here we evaluated how well preclinical models recapitulate human cancer and non-diseased tissue gene expression patterns *in silico* with respect to the full gene expression profile as well as subsetting by the most variable genes, genes significantly correlated with tumor purity, and tissue-specific genes by using publicly available gene expression profiles across multiple sources. We found that using the full gene set improves correlations between preclinical model and tissue global gene expression profiles, confirmed that GBM PDX global gene expression correlation to GBM tumor global gene expression outperforms GBM cell line to GBM tumor global gene expression correlations, and demonstrated that preclinical models in our study often failed to reproduce tissue-specific expression. While including additional genes for global gene expression comparison between cell lines and tissues decreases the overall correlation, it improves the relative rank between a cell line and its tissue of origin compared to other tissues. Our findings underscore the importance of using the full gene expression set measured when comparing preclinical models and tissues and confirm that tissue-specific patterns are better preserved in GBM PDX models than in GBM cell lines. Future studies can build on these findings to determine the specific pathways and gene sets recapitulated by particular preclinical models to facilitate model selection for a given study design or goal.

## 1. Introduction

In vitro preclinical models like cell lines and patient-derived xenografts (PDXs) are critical for advancing oncology studies but have limitations in how well they recapitulate human disease gene expression patterns and perform in the lab.(1) For example, cancer cell lines are generally less expensive to acquire and maintain, are more readily available from repositories, are more easily genetically manipulated, and have higher proliferation rates than PDXs.(2) However, because cancer cell lines are homogeneous, they fail to recapitulate disease and tissue heterogeneity and the microenvironment.(2) Additionally, over time, cancer cell lines are prone to mislabeling, cross-contamination, and contamination by chemicals (e.g., endotoxins, impure media) or biological entities (e.g., mycoplasma, bacteria, yeast).(3) In contrast, PDX models contain an *in vivo* microenvironment, are heterogenous, and have been shown generally to better match the patient’s tumor profile than cell lines.(4) In regards to both model types, the process and method of culturing can influence cell gene expression patterns.(5) As resources like the CelloSaurus and groups like the International Cell Line Authentication Committee continue to identify preclinical models that, through authentication testing, have been determined to be likely misidentified or contaminated, studies must continue to re-evaluate which preclinical models should be used in research.(6)

However, despite demonstrating efficacy in preclinical model systems like cancer cell lines and PDXs, most novel cancer drugs fail in clinical trials. Of all oncology drugs entering Phase I clinical trials, only 5% are approved. The main bottleneck of drug failure is in Phase II of clinical trials, suggesting that preclinical model systems are effective in predicting human drug toxicity (Phase I), but fail to accurately recapitulate many diseases for the sake of drug testing (Phase II).(7,8) Further, previous studies suggest that cancer cell lines often do not accurately represent the transcriptomic profile of their tissue or disease of origin, or that they may be better surrogates for particular cases, such as primary versus secondary tumors, or for a particular molecular pathway.(9,10) While disease and genetic background heterogeneity in humans and system or microenvironmental effects contribute to this, a better understanding of how cancer cell line or PDX model gene expression profiles recapitulate the human tumor or non-diseased tissue being modeled is critical.

Here we evaluated how restricting to the most variable genes impacts how well cancer cell lines or glioblastoma (GBM) PDX global gene expression profiles correlate with human tissue gene expression profiles. We used gene expression profiles from the Cancer Cell Line Encyclopedia (CCLE, n = 943),(11) the Human Protein Atlas (HPA; n = 194),(12) a publicly available osteosarcoma and bone sample set (SRP090849, n = 212),(13) GBM PDXs from the Mayo Clinic Brain Tumor PDX National Resource (n = 65),(14) The Cancer Genome Atlas (TCGA, n = 10,098)(15), and Genotype-Tissue Expression project (GTEx, n = 17,510)(16). We further investigated whether the inclusion or exclusion of tumor purity-correlated genes impacts those correlations and the ability of the preclinical models to capture tissue-specific gene expression signatures. Our findings illustrate that while GBM PDX samples outperformed GBM cancer cell lines concerning both tissue and disease context of origin, cell lines better model tissues of different origins, further underscoring that PDX models more faithfully capture tissue and disease-specific gene expression profiles. The degree to which either model captured the gene expression profile of the same tissue of origin, however, varied as expected depending on the gene set included when gene subsets were used. We find that it is generally best to assess global profiles, but if considering based on a smaller subset, like a specific pathway, it may be best to focus more consideration on this gene set of interest for informed decision making.

## 2. Methods

### 2.1 Data Collection

All analyses were conducted using the R programming language (version 4.2.2) within the RStudio interface (version 2022.02.01+461). We used the Recount3 R package to access publicly available RNA-seq gene counts (version 1.4.0)(17) GENCODE version 26 annotations (n=63,856 genes). All included data were originally generated using either an Illumina Genome Analyzer or HiSeq instrument. Data accessed through Recount3 was previously preprocessed with the Monorail pipeline, which facilitates comparison between cohorts and minimizes batch differences.(17) For this study, we used data from human tumor and non-diseased tissues, cell lines, and patient-derived xenograft (PDX) mouse models. Human tissue profiles were from The Cancer Genome Atlas (TCGA, n = 10,098 across 33 cancer types)(15), a publicly available osteosarcoma tissue data set (SRP090849; osteosarcoma n = 188, non-diseased bone n = 9)(13) to provide a bone tissue comparison for bone tissue-derived cell lines in subsequent analyses, the Genotype-Tissue Expression (GTEx) Project (n = 17,510 across 30 tissue types),(16) and the Human Protein Atlas (HPA, n = 171).(18) We sourced cell line gene expression data from the Cancer Cell Line Encyclopedia (CCLE, n = 1,004),(11) the HPA (n = 23),(18) and SRP090849 (bone-derived cell lines, n = 15).(13) In total, this data included 64 different cancers derived from 41 tissues. Finally, we sourced GBM PDX gene expression data from the Mayo Clinic’s Glioblastoma PDX National Resource (n = 65).(14)

### 2.2 Data Curation

We excluded some samples from subsequent analyses based on mislabeling or possible contamination (Supplemental Table 1). To avoid possibly confounding our analyses, we excluded the GTEx samples marked as bone marrow as they are derived from a leukemia cell line. Five HPA non-diseased tissue samples (ERR315357, ERR315383, ERR315386, ERR315403, and ERR315438) were relabeled as well for metadata consistency. We selected primary tumor samples within the TCGA dataset. We also confirmed that each CCLE cell line had not been flagged as having evidence of contamination or misidentification in The Cellosaurus, a Swiss Institute of Bioinformatics resource for biomedical cell lines.(6) From this assessment, we removed 66 of the CCLE cell lines from our analysis,(6) as well as one additional sample (SRR8615727) which was missing significant portions of its metadata, including which cell line it was derived from. We also relabeled 4 HPA cell lines (SRR629581, SRR629582, SRR4098609, and SRR4098610) for metadata consistency in subsequent analyses. We removed one PDX sample (SRR9294079) as it is IDH-mutant and no longer classified as GBM by the World Health Organization as of 2021.(19)

### 2.3 Data Categorization

In order to match preclinical models to corresponding tissues for cross-disease and cross-tissue comparisons we assigned preclinical models by their putative tissue of origin (Supplemental Tables 2 and 3). For example, all leukemia-derived cell lines regardless of subtype were grouped as “Blood Cancers”. With these adjustments, all cell lines were identified as part of a parent tissue group. However, some supplementary tissues (i.e., those without corresponding preclinical models in our analyses) were also included to further evaluate model performance in contexts separate from their origins (e.g., GBM cell lines and PDXs evaluated for their correlation to other non-brain tumor tissue types).

### 2.4 Data Normalization

From raw counts, we calculated transcripts per million (TPM) using the gene lengths available in the Recount3-downloaded ranged-summarized experiment (RSE) object with merged Ensembl/Havana GENCODE version 26 annotations(20) via the getTPM() function of the recount R package (version 1.20.0) to facilitate a comparison of the proportion of total reads mapped to a gene across samples.(21)

### 2.5 Gene Subsets

Previous studies evaluating gene expression differences between cell lines and tissues by restricting to the 5,000 most variable genes(9,10,22,23) suggest this subset represents the most “likely biologically informative”(9,10) or representative signal.(22,23) This restriction of the global gene set may be useful for reducing computational memory requirements and run times, but research is limited on biological information gained or lost with varying gene subsets. We evaluated the difference in variability across samples by gene set size with principal component analysis (PCA) with the mixOmics R package (version 6.18.1) pca() function to determine principal components,(24) and then used the PLSDAbatch package (version 0.2.1) Scatter_Density() function to generate PCA and density rugplots to investigate the variance between sample origin types (i.e., cell line, PDX, non-disease tissue, tumor tissue; between brain-derived tissues and GBM-derived models in Figure 2B; between all samples of all sources in Supplemental Figure 1A), as well as with respect to other variables that may impact variance (i.e., read length, tissue type, sample source, sex; Supplemental Figure 1B-E).(25)

We identified tumor purity-correlated genes based on TCGA tumor gene expression significantly correlated with ABSOLUTE-generated tumor purity. We also defined tissue-specific gene expression from GTEx using the Harminozome (accessed November 2022).(16,26,27) We chose this resource due to its breadth of tissue-specific gene sets available across various tissues.

### 2.6 Correlation & Pathway Analysis

We used Spearman’s rank correlation with the cor() function of the stats base R package(28) to evaluate model performance in recapitulating tissue gene expression patterns (Supplemental Table 4). We visualized the correlation distributions with the hist() function in the graphics base R package (GBM cell lines correlated to brain tumor tissue: Figure 5A; all other groups: Supplemental Figure 2).(28) We computed the significance between correlation performance using the Wilcoxon rank sum test using the rstatix (version 0.7.0) and coin (version 1.4-2) R packages.(29–31) We studied their performance within many sub-groups. To investigate pathway enrichment of genes highly and lowly correlated and anticorrelated between brain-derived models and brain tumor and non-diseased tissue used the gprofiler2 R package (GOSt results for these gene groups are in Supplemental Table 5).(32) We generated visualization of correlations, pathways, and all other figures using the ggplot2 (version 3.3.5), ggpubr (version 0.4.0), ComplexHeatmap (version 2.10.0), mixOmics (version 6.18.1), and PLSDAbatch (version 0.2.1) R packages and compiled with BioRender.(24,33–37)

## 3. Results

### 3.1 The correlation between cancer cell line and tissue of origin gene expression when restricting to the most variable genes is reduced and less specific compared to the full gene expression profiles

Previous studies evaluating gene expression differences between cell lines and tissues by restricting to the 5,000 most variable genes(9,10,22) suggest this subset represents the most biologically informative signal.(10) We evaluated how the gene subset size impacts the correlation between cancer cell line and tumor tissue gene expression profiles originating from the same tissue type (i.e., with matched origin tissue). From CCLE,(11) HPA,(12) and SRP090849(13) cancer cell line gene expression profiles, we calculated their global Spearman correlation to the corresponding non-diseased tissue (GTEx,(16) HPA, SRP090849) or tumor tissue profiles (TCGA,(15) SRP090849). For this analysis, we determined the 100, 1,000, 5,000, and 10,000 most variable genes across all included TCGA tumor tissue samples or GTEx non-diseased tissue samples. We then evaluated how restricting each of those subsets of genes, impacted the correlation between cancer cell lines and matched tissue types compared to the global profile (n = 63,856 genes) (Figure 1A). We also evaluated including or excluding previously identified high tumor purity-correlated genes (i.e., TCGA tumor gene expression significantly correlated with ABSOLUTE-generated tumor purity).(10,38,39)

**Figure 1.**
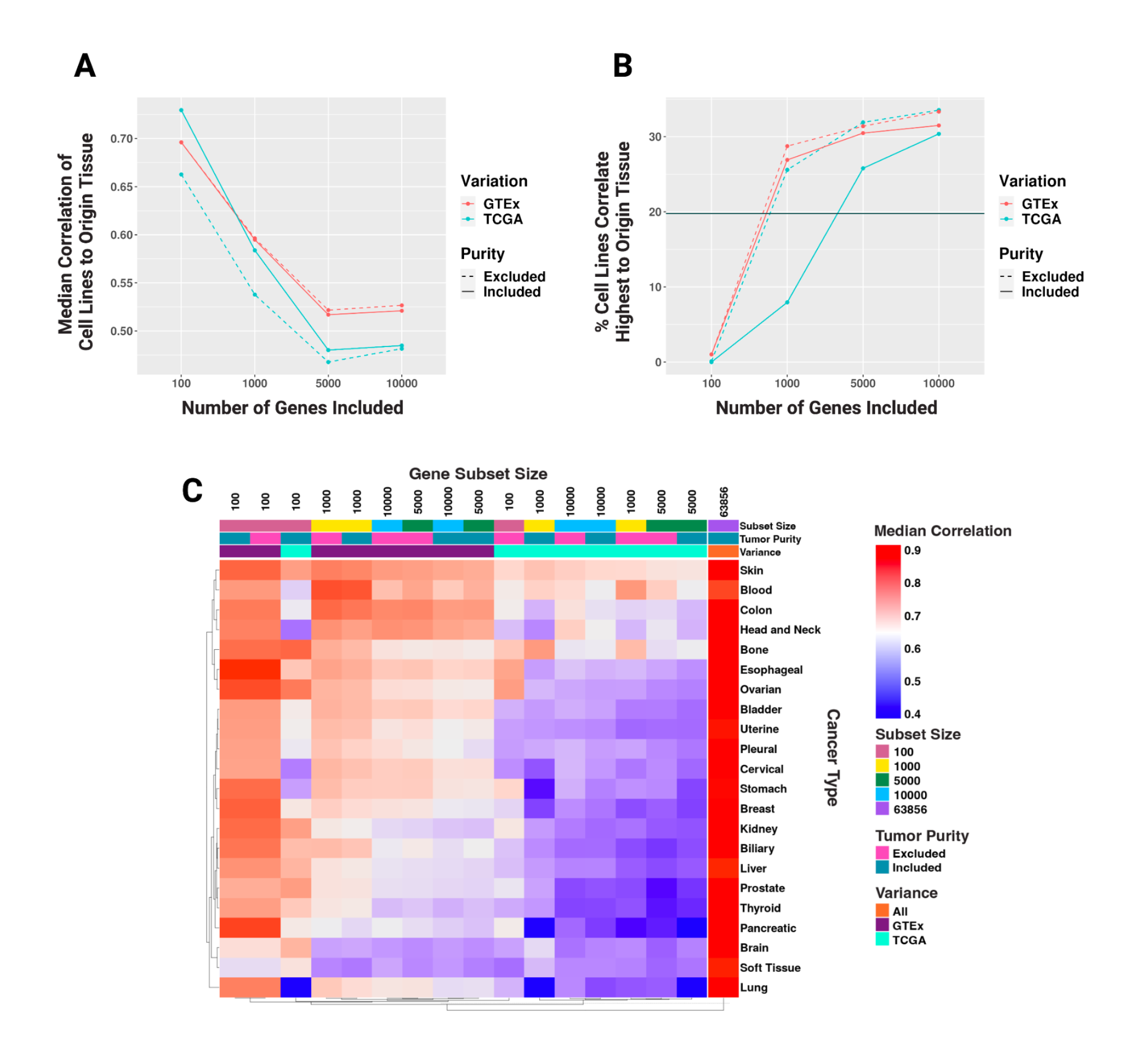
**A.** Line graph of the difference in median correlation of cell lines to their matched origin tumor tissue based on the set size of most variable genes, whether that variation is based on GTEx (red) or TCGA (blue) most variable gene subsetting, and whether tumor purity-correlated genes are included (solid line) or excluded (dashed line). Global correlation is represented as a gray solid line. **B**. Percent of cell lines that correlate highest with their matched origin tumor tissue out of all tumor and non-diseased tissues included based on the set size of the most variable genes, whether that variation is based on GTEx (red) or TCGA (blue) tissue variation, and whether tumor purity-correlated genes are included (solid line) or excluded (dashed line). Global correlation is represented as a gray solid line. **C**. Heatmap of cell line median correlation to their matched origin tumor tissue based on the set size of most variable genes, whether that variation is based on GTEx or TCGA tissue variation, and whether tumor purity-correlated genes are included.

Compared to the median correlation between the cancer cell lines and their matched tumor tissue in the global gene profile (rho = 0.82) (Supplemental Table 4), the 100 most variable gene subset based on TCGA variance, including tumor purity-associated genes, had the highest median correlation value (rho = 0.73).

However, the 5,000 most variable gene subset based on TCGA variation, excluding tumor purity-associated genes, had the lowest correlation value (rho = 0.47). The median correlation rho between cancer cell lines and matched tumor tissue in the 5,000 most variable genes excluding tumor purity-correlated genes, was 0.05 greater in GTEx-derived variation than in TCGA. Therefore, we found that restricting to the most variable genes reduces the cell line-to-tissue of origin correlation and that cell lines generally correlated more to the same tissue they were derived from when using the most variable genes based on GTEx samples. Further, while excluding purity-associated genes improved the correlation for gene sets based on the most variable GTEx genes, it did not for the most variable genes across TCGA samples (Figure 1A). As expected, the purity-associated genes were the most correlated between cell lines and tumor samples due to the non-tumor cells within the sample.(39)

We also evaluated the percentage of cancer cell lines that correlated most highly with their own matched tissue of origin compared to all other tissues included in this study (i.e., both cancer- and non-disease-derived tissues). Intriguingly, even though cancer cell line model gene expression profiles were most strongly correlated with their tissue of origin when only the top 100 variable genes were included (Figure 1A), very few correlated most highly with their tissue of origin when compared to any tissue (i.e., 0% and 0.1% when the variable genes were selected based on TCGA and tumor purity-correlated genes were included and excluded, respectively; 1% when the variable genes were selected based on GTEx, regardless of the inclusion of tumor purity-correlated genes). Further, while 20% of cell lines correlated most highly to tumor tissues of the same origin with the global gene expression profile, that percentage was higher in all gene subsets greater than 100 except for in the 1,000 genes based on TCGA variation with tumor purity-correlated genes included (Figure 1B). That is to say, while cell lines had higher correlation rho values within smaller subsets of top varying genes, they had less specificity to their origin tissue.

We further investigated the effect of these different gene subsets on the correlation of cell lines to tissues at the individual tissue level, to see if these trends were consistent across tissues or tissue-specific (Figure 1C). We observed that some tissue types had less variation in their global gene expression correlation to cancer cell lines from that same tissue type regardless of gene subset size, GTEx or TCGA tissue gene expression most variable gene subsetting, or purity gene inclusion. That is, certain tissues consistently correlated highly (e.g., skin) or lowly (e.g., brain) with cell lines from their matched origin regardless of subset.

For example, when comparing most variable genes based on GTEx or TCGA, the difference between the median correlation of all of the most variable gene set correlations (i.e., all subset sizes, tumor purity gene inclusion/exclusion) was ∼0.01 for soft tissue cancer-derived tissue (i.e., soft tissue/connective tissue cancer, sarcomas; rho values ranged between 0.53 and 0.67) and ∼0.02 for bone and brain cancer-derived tissue (rho values ranged between 0.64-0.79 and 0.53-0.71, respectively). However, the difference in rho was at least 0.1 in 14 of the 22 tumor tissue types we analyzed. This suggests that for many tissues, how the most variable genes are defined for the purpose of comparing cancer cell lines to tumor tissues from the same tissue of origin impacts the results. Based on these findings, for subsequent analyses, we included all genes.

### 3.2 Global gene correlations between GBM models and brain tissues underscore GBM PDX gene expression specificity and GBM cell line gene expression ambiguity

We next correlated GBM cell line and PDX gene expression profiles to brain cancer and non-diseased brain tissue profiles. Principal component analysis (PCA) demonstrated that principal component 1 (39% of the variance) and principal component 2 (15% of the variance) separated non-diseased brain tissue from GBM PDXs, GBM cell lines, and brain tumor tissue (Figure 2A). To evaluate how well the GBM PDXs and cell lines recapitulated GBM-specific, brain tumor-specific (i.e., GBM and lower-grade glioma, LGG), and brain non-diseased tissue gene expression profiles, we compared the median correlation rho values for each (Figure 2B). We found that GBM PDX models best correlated to brain tumor tissues, and both GBM cell lines and PDX models correlated better to tumor tissue than non-diseased tissue. We found that, though within a modest correlation range (all samples of both models’ median correlations to either tissue type were between Spearman rho = 0.77-0.89) PDXs significantly correlated more to brain tissue, both tumor and non-diseased, than cell lines (p < 0.0001 and W = 410 when correlated to brain tumor tissue, p < 0.0001 and W = 455 when correlated to brain non-diseased tissue, Figure 2C). Finally, we asked how well GBM PDX and cell line global gene expression profiles correlated to other tumor and non-diseased tissues from TCGA and GTEx because other studies have suggested cell line gene expression profiles may better resemble other tumor types than the one from which they were originally derived.(40) Here, we found that the GBM cell lines correlated significantly better to non-brain tissues than the GBM PDX gene expression profiles did (Bonferroni-corrected Wilcox p-value < 0.05, Figure 2D) in every comparison with the exception of two tumor tissues (i.e., adrenal and eye), and five non-diseased tissues (i.e., pituitary, colon, nerve, testis, and ovary). This suggests that the GBM PDXs better recapitulate their tissue of origin global gene expression patterns and that the GBM cell line global gene expression profiles are less tissue-specific and therefore correlate better with both cancer or non-diseased tissues different from their own origin than the PDX models.

**Figure 2.**
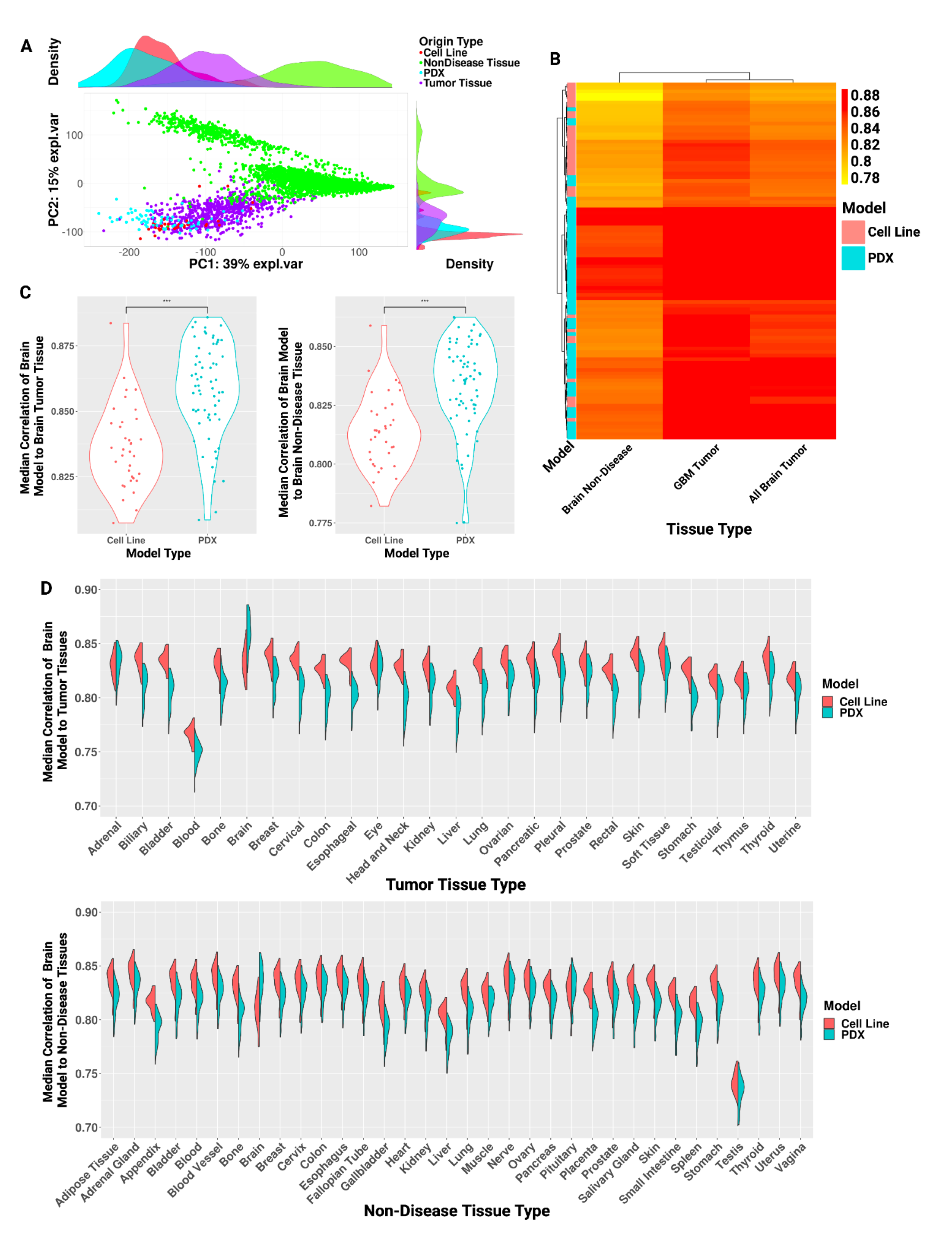
**A.** Principal component analysis scatterplot with density rug plot of gene expression profiles from GBM cell lines (red), GBM PDX models (teal), non-diseased brain tissue (green), and GBM and LGG tissue samples (purple). **B**. Heatmap of median performance of each GBM cell line and PDX sample to brain non-disease, GBM tumor, and all available brain tumor (GBM, LGG) tissue. **C**. Violin plot showing median correlation of each GBM cell line and PDX sample to brain tumor (left) and non-diseased (right) tissue split by model type. Significant Wilcox p-value (p < 0.05) indicated by asterisks. **D**. Split violin plot of correlation of each GBM cell line and PDX sample to the tumor (top) and non-diseased (bottom) tissues. Wilcoxon rank sum test with Bonferroni procedure for multiple hypothesis correction, p-adjusted values < 0.05 indicated by asterisks and colored by model type with significantly greater median correlation.

### 3.3 Models often fail to recapitulate origin tissue expression of tissue-specific genes

Given the importance of tissue specificity to model gene expression fidelity, we further considered how well cell lines recapitulated their matched tissue of origin (in non-diseased and tumor contexts) when subsetting gene expression profiles to tissue-specific genes for each respective tissue. Here we defined genes as having tissue-specific gene expression based on prior work identifying genes with either high or low gene expression in each tissue relative to all other tissues in the GTEx dataset.(16,27) For each tissue, we calculated the correlation between the gene expression of that tissue’s tissue-specific genes in cancer cell lines derived from that tissue to the non-diseased gene expression profiles of that tissue and cancer gene expression profiles of that tissue (Figure 3A). In 14 out of 16 tissues, the tissue-specific gene expression correlation is higher between cancer cell lines and tumor tissues than between cancer cell lines and non-diseased tissue for that tissue (Bonferroni-corrected Wilcoxon p < 0.05).

**Figure 3.**
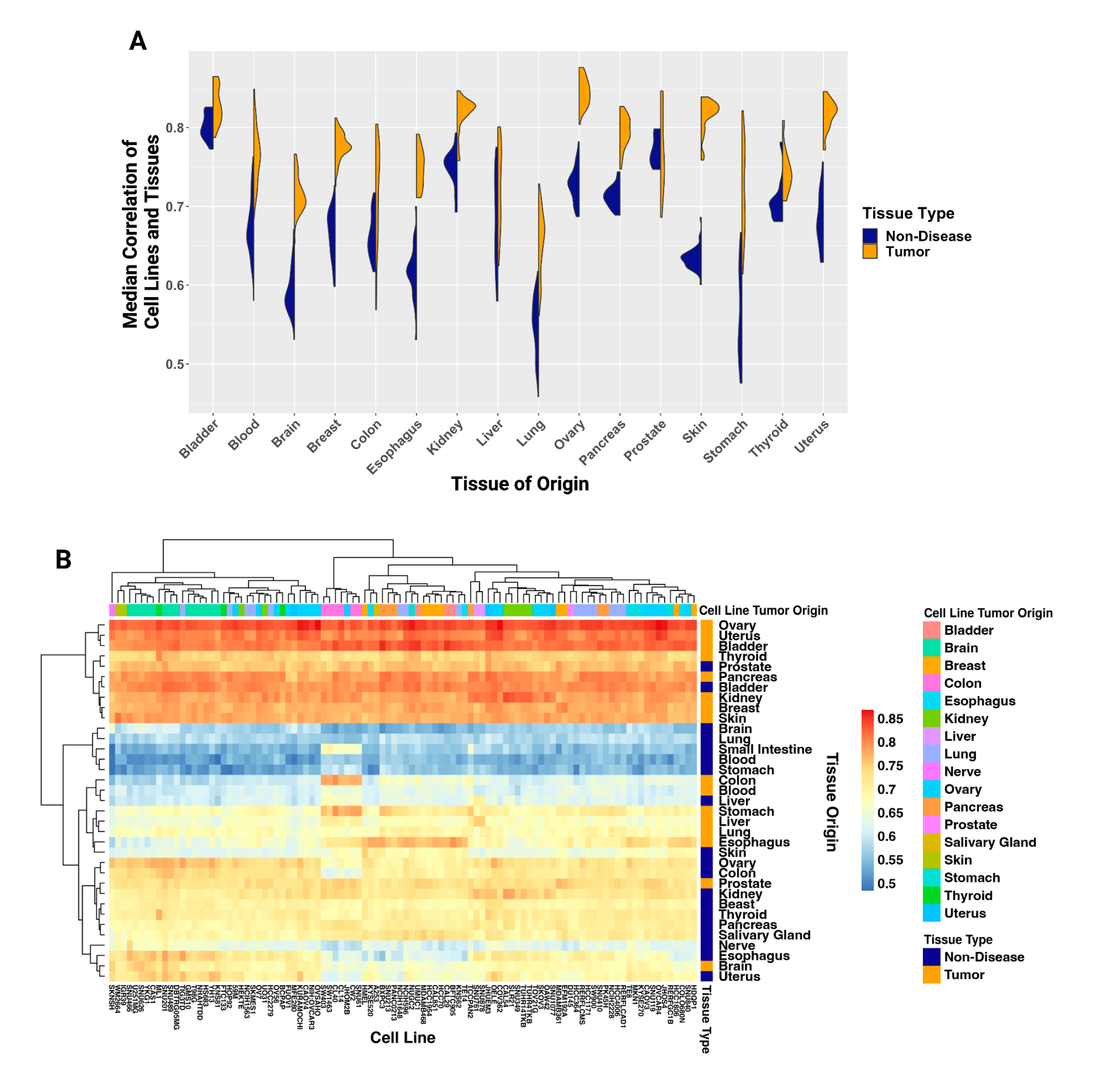
**A.** Violin plots of the median correlation of each cell line sample to its matched tumor (orange) and non-diseased (dark blue) tissue in each matched tissue’s specific genes. Wilcoxon rank sum test with Bonferroni procedure for multiple hypothesis correction, p-adjusted values < 0.05 indicated by asterisks and colored by tissue type with significantly greater median correlation. **B**. Heatmap of correlation of the top 100 most generally highly correlated cell lines to each tumor and non-diseased tissue by tissue-specific genes.

We next studied the performance of the 100 cancer cell lines with the highest median global gene expression correlation to all studied tissue and disease contexts with respect to a given tissue’s tissue-specific genes. When visualized as a heatmap of the cancer cell lines’ correlation to each tissue and disease context for that tissue’s tissue-specific genes (Figure 3B), we found there were subclades of some cancer cell lines from the same tissue of origin clustering together (e.g., some brain-derived and liver-derived cell lines), but, as noted in previous studies, this was not widespread.(41) Some tissues clustered by tumor or non-diseased state, though again not all (e.g., within the first split of the dendrogram, eight of the ten tissues were tumor-derived, but the other two were non-diseased prostate and bladder). Though cell lines included here were derived from 25 different tissues, all of the top 100 cancer cell lines, regardless of origin, correlated highly (rho > 0.72) with kidney, breast, skin, thyroid, ovary, pancreas, uterus, and bladder tumor tissue as well as prostate and bladder non-diseased tissue. Further, all included cancer cell lines correlated less with brain, lung, blood, and stomach non-diseased tissue (rho < 0.66). As cancer cell lines were more correlated on average to tumor gene expression profiles, this further underscores that tissue status (i.e., from a tumor or non-diseased tissue) is a critical driver of how well a cancer cell line’s global expression profile correlates with that tissue.

These tissues had low variation in the strength of correlation across cell lines (e.g., bladder non-diseased tissue had a difference of 0.04 between the lowest and highest correlation rhos); however, a few tissues, such as colon tumor and non-diseased esophagus, had higher variation in how well they correlated with the 100 tested cancer cell lines (i.e., the difference between lowest and highest correlation rho between cell line and tissue for colon tumor and non-diseased esophagus were 0.25 and 0.22, respectively). Further, 18 of 35 tissues had a difference between their lowest and highest rho correlation value of at least 0.1. Colon tumor samples predominantly correlated highly with colon-derived cell lines; all 6 of the colon-derived cell lines correlated highest to colon tumor tissue (rho > 0.7). However, other tissues, including pancreatic tumor tissue, had less specificity and correlated highly with cell lines across origins. Only 3 of the 6 pancreatic tumor-derived cell lines included were in the top 15 rho values of pancreas tumor tissue’s correlation to cell lines, while the lowest correlation between pancreas tumor tissue and pancreas tumor-derived cell line was the 36th highest rho value at rho = ∼0.81, and no correlation to pancreas tumor tissue out of any of the 100 cell lines was lower than rho = ∼0.75. Ovary tumor tissue correlated highly with all of the top 100 cell lines (the lowest rho was = ∼0.82), and with minimal specificity to cell lines of the same origin; while 10 of the 19 ovary cell lines were in the top 15 rho values of ovarian tumor tissue’s correlation to cell lines, one of those 19 was ranked 95th for its strength of correlation (rho = ∼0.825). This supports that some tissues are more broadly recapitulated by cancer cell lines regardless of the cell lines’ origin, while others have greater specificity, whether that be to cell lines derived from the same matched tissue or a specific set of tissues.

We again focused on GBM cell lines and PDXs and found that while GBM PDXs were more correlated to both brain-derived tumor and non-diseased brain tissue than GBM cell lines in the global profile (Figure 2C) when restricted to tissue-specific genes, this difference was even greater (median Spearman correlation rho of PDX gene expression to tumor gene expression and to non-diseased brain tissue gene expression was 0.126 and 0.155 greater than to the GBM cell lines, respectively) (Figure 4A). This difference is mainly due to GBM cell lines having a reduced correlation to brain tumors and brain tissues when subsetting for brain-specific genes rather than full gene profiles, similar to what we previously found with varying gene set sizes (Figure 1A). We then asked how our GBM models correlated to non-brain tumors and tissues based on each tissues’ tissue-specific genes to assess how specific models are to their origin disease and tissue contexts (correlation to tumor tissue: Figure 4B; correlation to control tissue: Supplemental Figure 2). We again found that brain-derived cancer cell lines better recapitulated gene expression of other tissues compared to GBM PDX models, but there was more variability here than compared to the full global gene expression profile. The range of median correlation to each tumor tissue for GBM cancer cell lines was 0.54-0.85 and for GBM PDXs was 0.46-0.88 using tissue-specific genes (Figure 4B), while in the full gene set median correlation for cancer cell lines was 0.75-0.86 and PDXs was 0.71-0.89 (Figure 2D). We concluded that PDX models’ ability to recapitulate tissues’ gene expression may be more origin context-specific (i.e., origin disease and tissue), while cell lines’ ability to recapitulate tissues’gene expression may be more ambiguous, recapitulating across non-origin tissues and correlating higher when not considering tissue-specific genes vs. global gene profiles.

**Figure 4.**
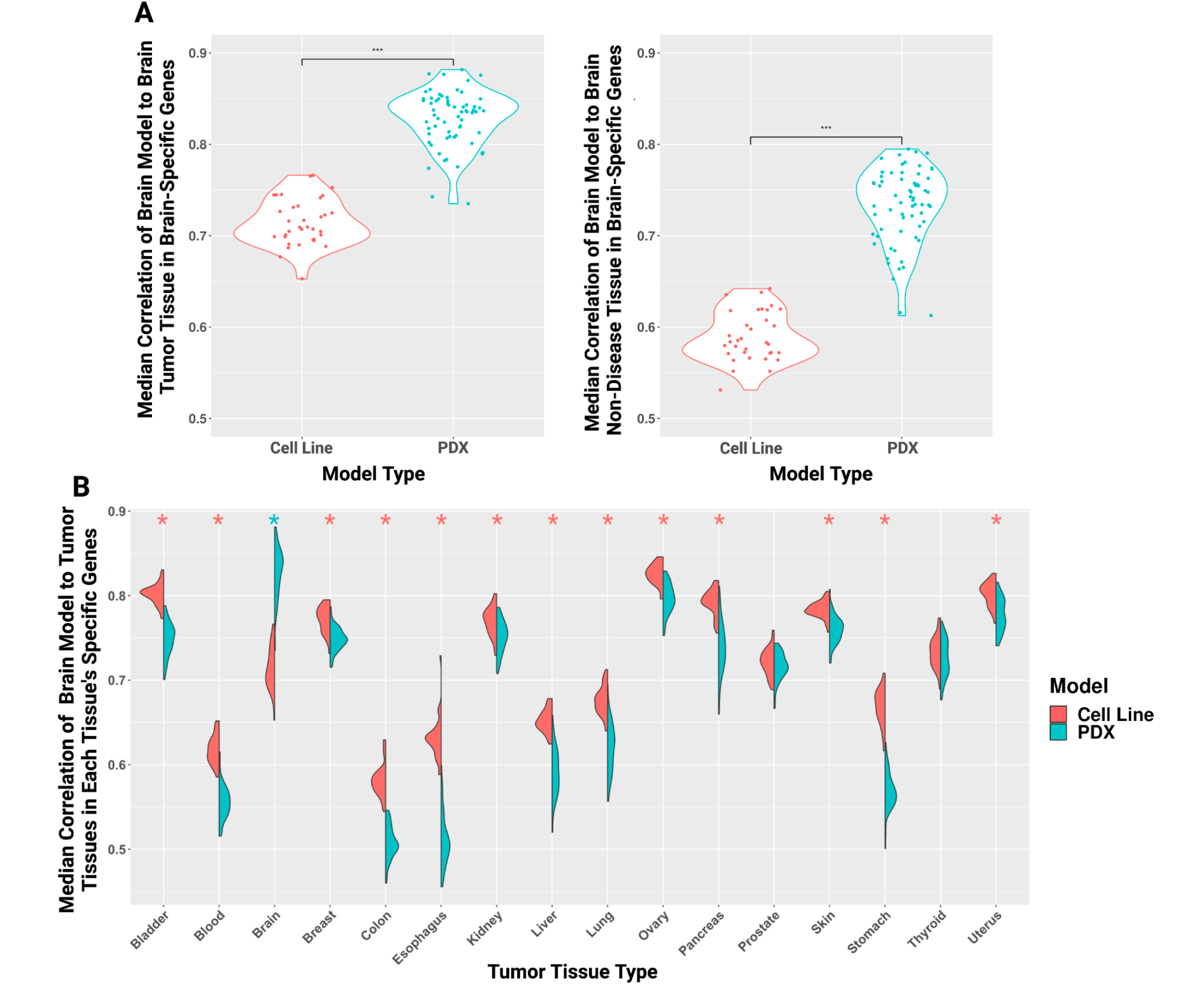
**A.** Violin plots with Wilcoxon test showing the median correlation of each GBM cancer cell line and PDX sample to brain tumor and non-diseased brain tissue with brain-specific genes. Significant Wilcox p-value (p < 0.05) indicated by asterisks. **B**. Split violin plots of the correlation of each GBM cancer cell line and PDX sample to each tumor tissue type by that tissue’s specific genes. Wilcoxon rank sum test with Bonferroni procedure for multiple hypothesis correction, p-adjusted values < 0.05 indicated by asterisks and colored by model type with significantly greater median correlation.

### 3.4 Gene ontology analysis of genes significantly correlated and anti-correlated between brain-derived models and brain tissue gene expression

Having assessed the gene expression correlation of cell lines and PDXs to matched tumors and tissues by global profiles and multiple subsets of genes, we then asked which genes in particular are successfully or unsuccessfully recapitulated by included GBM-derived models in comparison to brain tumor or non-diseased tissue, and if these genes are enriched for any pathway or biological terms. To do this, we determined which genes were highly or lowly correlated or anticorrelated between GBM models and brain tumor or non-disease tissue gene expression profiles (Figure 5A; Supplemental Figure 3), and if any terms were overrepresented within these sets using g:Profiler (Supplemental Figure 4; Supplemental Table 5).(42) We found that most genes significantly correlated between GBM cell lines and brain tumor or non-diseased tissues did not overlap with genes significantly correlated between GBM PDXs and brain tumor or non-diseased tissues (Figure 5B; Supplemental Table 5). We found some of the genes significantly correlated between GBM models that were exclusive to either PDXs or cell lines were transcription factors, including *DACH2, ELK3, FEV, GTF2H2B, KLF10, MAFK, REST, SALL4P5*, and *TFB1M*, specific to PDXs and *HSFX4, HSFY8P, MITF, MYCN, SP2*, and *YY2*, specific to cell lines. The top terms significantly enriched for each group were unique to a given comparison (Figure 5C; Supplemental Table 5). When evaluating between GBM cell lines and brain tumor tissue, we found that cellular response to heat shock terms were overrepresented in correlated genes, while terms related to immune response (e.g., NF-kappaB complex, Innate Immune System, Neutrophil degranulation) were overrepresented in anticorrelated genes (Figure 5C; Supplemental Table 5). Between GBM cell lines and non-diseased brain tissue gene expression, highly correlated genes were, surprisingly, overrepresented for oxidative phosphorylation and related terms (e.g., respiratory chain complex), and anti-correlated to cellular and anatomical development and cellular motility (Figure 5C; Supplemental Table 5). The terms significantly enriched in the correlated gene set between GBM PDXs and tumor brain tissue varied widely, from Bardet-Biedl syndrome, microtubule cytoskeleton, “skin 1; Langerhans”, to exercise-induced circadian regulation. The anticorrelated gene set had top enriched terms for oxidoreductase activity and metabolic pathways (daunorubicin metabolic pathway, polyketide metabolic process, etc.) (Figure 5C; Supplemental Table 5). Gene sets significantly correlated between non-diseased brain tissue and the GBM PDXs included cell-cell signaling and synaptic signaling. The terms from the significantly anticorrelated gene set included mostly metabolic pathways (e.g., cholesterol biosynthesis, alcohol biosynthetic process, and steroid biosynthesis). Our results here suggest that cell lines and PDX models recapitulate disease contexts in unique biological terms and pathways, and, further, uniquely fail to capture other specific biological terms and pathways.

**Figure 5.**
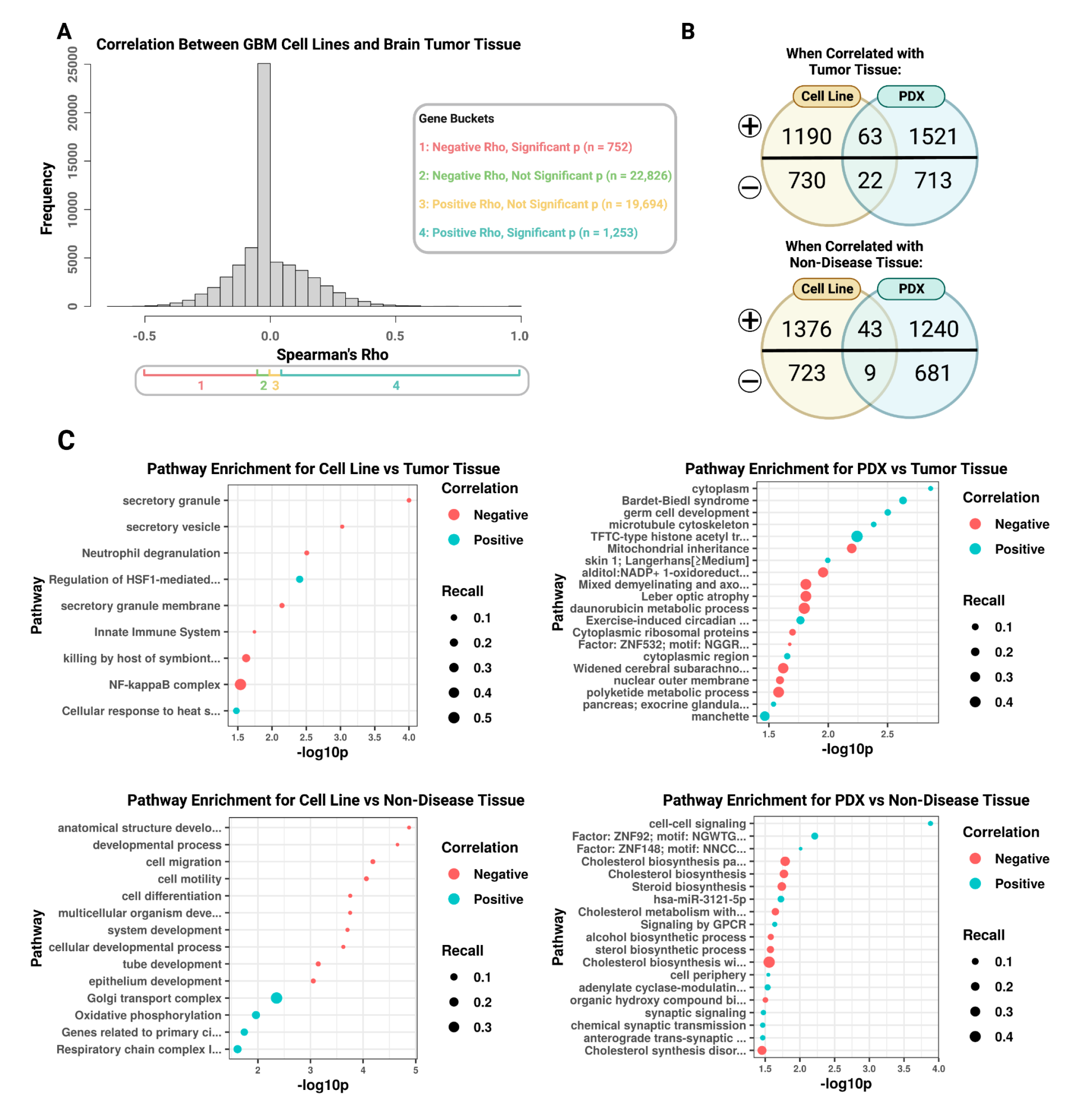
**A.** Histogram of the distribution of Spearman’s correlation between GBM cell lines and brain tumor tissue and the separation into groups by positive rho (i.e., correlated) or negative rho (i.e., anticorrelated) as well as the p-value (i.e., p ≥ 0.05 is insignificant, p < 0.05 is significant). **B**. Venn diagrams with the number of genes significantly correlated (top of each diagram, denoted by “+”) or anti-correlated (bottom of each diagram, denoted by “-”) between GBM models (cell lines on the left, PDXs on the right) and brain tumor (top diagram) and non-diseased (bottom diagram) tissue. **C**. Dot plots of the most significant (top 10 if at least 10 were significant) pathways represented in significant genes with either positive (blue) or negative (red) correlation between, clockwise from top left, GBM cell lines and brain tumor tissue, GBM PDXs and brain tumor tissue, GBM PDXs and brain non-diseased tissue, and GBM cell lines and brain non-diseased tissue, respectively. Dots are sized by recall or the proportion of genes from a specific term represented in the data divided by the total genes possible for that term.(43)

## 4. Discussion

In this study, we evaluated how well cancer cell lines derived from many different tissues recapitulate human tissue expression patterns across different contexts (i.e., tissue, disease) and concerning gene inclusion (i.e., most variable genes, purity associated genes, tissue-specific genes). We also performed additional analyses in GBM models, given the availability of GBM PDX gene expression profiles. We found that using the full gene expression profile improves correlations between preclinical model and tissue global gene expression profiles, confirmed that GBM PDX global gene expression correlation to GBM tumor global gene expression outperforms GBM cell line to GBM tumor global gene expression correlations, and demonstrated that preclinical models in our study often failed to reproduce tissue-specific expression. While including additional genes for global gene expression comparison between cell lines and tissues decreases the overall correlation, it improves the relative rank between a cell line and its tissue of origin compared to other tissues.

Previous studies have sought to prioritize cell lines for disease recapitulation by correlating the tumor type of interest to cell lines by the 5,000 most varying genes across samples, typically reasoning that this subset of genes are likely to be biologically informative.(9,10,22) For this reason, we investigated how correlation values change across gene subsets of various sizes and how they compared to using the entire gene set (Figure 1). Further, we are not aware of any similar studies that have attempted to prioritize cancer cell lines by the fidelity of the gene expression profile compared to non-diseased tissues. However, we suggest that comparison of cell lines to non-diseased tissues is important for selection of preclinical models that may better recapitulate non-cancer disease contexts, considering genetic drift and other abnormalities of tumors.(44,45) We found that by using subsets of the most varying genes (i.e., 100, 1,000, 5,000, and 10,000), cell lines generally correlated less to their tissue of origin (both tumor and non-diseased), and yet that the full gene sets correlated higher than any varying gene subset (median correlation rho = 0.82) (Figure 1A).

Celligner (previously developed by the DepMap team) is a method that uses an unsupervised alignment approach to compare protein-coding gene expression of cell lines to tumor tissues to correct for differences in tumor compared to cell lines. They found that cell lines matched closest to their disease of origin (e.g. kidney tumor-derived cell line matching best to their kidney tumor cluster) 57% of the time.(40) Other studies correlating gene expression of cell lines to tumors by the 5,000 most varying genes found only 8 out of 22 tumors correlated highest with cell lines derived from the same tumor type.(10) Similarly, we found only ∼32% of cell lines matched first to their disease/tissue of origin (Figure 1B). However, the proportion of cell lines matching best to their disease/tissue of origin increased with increasing varying gene subset size, with the exception of the full gene set, which only correlated higher than the top 100 varying gene subset. This is surprising compared to the trend we saw when comparing median correlation values of cell lines to their origin tissues, and shows that the highest correlation may not necessarily show the best recapitulation of the disease/tissue it was derived from (e.g., while the top 100 varying genes were most highly correlated between a cell line and its disease origin tissue, it may still correlate higher with a different disease/tissue). We further considered the specific tissues and compared the correlations of these gene subsets by each tissue and disease context, finding that the impact of gene subset on correlation varied by tissue (Figure 1C). Certain tissues consistently correlated highly across subsets (e.g., skin), while others consistently correlated lowly across subsets (e.g., brain). However, other tissues’ correlation rho values varied greatly to matched cell lines by subset (i.e., the most varying gene subset size, tumor purity inclusion/exclusion, etc.) Consistent with previous studies, we found pancreatic and lung cancer comparisons to have some of the most variable correlation coefficients across cell lines, believed to be due to greater heterogeneity within those tumor types.(10)

Similarly to previous findings in neuroblastoma, our comparisons of GBM PDX models and GBM cell lines supported that PDX models were significantly more similar than cell lines to their matched disease context (Figure 2).(46) This is unsurprising, as the PDX model have a microenvironment, vascularization, and cellular heterogeneity, all critical for recapitulating human tissue.(2,4) However, our study is the first to our knowledge to further compare PDX and cell line models to a non-disease tissue context. As previously mentioned, preclinical model recapitulation of non-diseased tissue for non-cancer contexts (e.g., rare diseases) is likely more informative than tumor recapitulation, due to multiple genomic abnormalities in malignancies.(44,45) When subsetting only GBM tumor-derived cell lines and PDXs, we found PDX models and cell lines both correlate highest to GBM-specific tumor tissue, followed by all other brain tumor tissue, and then non-diseased brain tissue (Figure 2B). While PDX model profiles correlated significantly higher to brain tissue regardless of disease context (p < 0.0001, W = 410-455), the range of correlations for specific models overlapped between PDX and cell lines (rho 0.77-0.89). As found in previous studies and here, cell lines often exhibit greater transcriptome expression similarity to other tissues than the one from which they were originally derived.(40) With this in mind, we compared GBM-derived cell lines and PDXs across other tumor and non-diseased tissue contexts and found GBM cell lines do significantly correlate higher with other tissue profiles than GBM PDXs across all contexts with the exception of two tumor tissues (adrenal and eye), and five non-diseased tissues (pituitary, colon, nerve, testis, and ovary) (Figure 2D). Interestingly, the only tissues that GBM PDXs correlated with higher than GBM cell lines did (though not significantl) were adrenal tumors and non-diseased pituitary.

To further investigate tissue-specific recapitulation of cell line gene expression profiles across disease contexts, we procured lists of tissue-specific genes and compared the correlation of cell line gene expression profiles to their matched tumor and non-diseased tissues based on expression of that tissue’s tissue-specific gene set.(16,27) This revealed 14 out of 16 tumor tissues had significantly higher correlation to their matched cell lines compared to their non-diseased tissue (Figure 3A). We found the 100 cell lines with the highest median correlation values by tissue-specific genes clustered by tissue origin for only a few tissue origins (i.e., cell lines derived from brain mostly clustered together as well as lung, Figure 3B). Further, tissues mostly clustered by disease context (tumors vs. non-diseased), particularly one subclade consisted of only non-diseased tissue (i.e., brain, lung, small intestine, blood, and stomach). This particular non-diseased tissue subclade also revealed the lowest correlation rho values, suggesting these tissues may be the most challenging to recapitulate gene expression profiles in a cell culture model, based on their tissue-specific genes. Indeed, others have demonstrated the challenge of modeling the tissue context accurately with cell lines in these tissues, leading to recommendations for patient primary derived cells and organoids for in vitro studies.(27,44,45,47–49)

We further compared our list of top 100 cell lines by median correlation to previous work by Yu et al. 2019, where they selected the five cell lines with highest correlation for each tumor type studied (110 total cell lines, 22 tumors) and found an agreement of 24 cell lines (Supplementary Table 4).(10) Cell lines unique to each study are likely due to differences in the gene subset used for calculating correlation (5,000 most varying protein-coding genes vs. tissue-specific genes) and then selection of top five cell lines median correlation for each tissue compared to selection of top 100 median correlation (regardless of tissue). This also underscores the importance of the gene subset selection for studies. Further, when subsetting for brain-specific genes rather than full gene sets, we observed a decrease in the correlation between GBM-derived cell line compared to brain tumor gene expression profiles, brain non-diseased tissue gene expression profiles, and across tissue gene expression profiles (Figure 2, Figure 4). Correlation values between GBM-derived PDXs and matched tumors or tissues were very similar when restricting to brain-specific genes versus full gene sets (Figure 2C, Figure 4A). However, when comparing across other tissues using their tissue-specific genes, GBM-derived PDXs had lower gene expression correlations than with the full gene expression profiles (Figure 2D, Figure 4B). This suggests that while tissue-specificity is a driver of the gene expression profile correlation between PDXs and tissues, it is not for cell lines.

When comparing significantly correlated genes between GBM preclinical models and brain tissue or tumor, we found most significantly correlated genes were exclusive to GBM PDX or GBM cell line to tissue correlations (Figure 4B). The expression of these genes may confer a better fit of one model type or the other, depending on the area of research. In particular, we highlight the transcription factors *FEV, MAFK*, and *REST*, whose expression was significantly positively correlated between tissues and GBM PDX models but not between tissues and GBM cell lines. *REST* expression has been previously found to be correlated with immune cell infiltration and immune checkpoints in glioma, and its expression has been recommended suggested as a biomarker of poor prognosis in glioma.(50) *FEV* is believed to regulate genes involved in early brain development and the serotonergic pathway, and *MAFK* is associated with numerous neurological disorders through oxidative stress response.(51,52) In contrast, a transcription factor whose expression was significantly correlated between brain tissues and GBM cell lines but not between brain tissues and GBM PDXs was *MYCN. MYC* and *MYCN* have been found to have specific and different Arf suppression mechanisms and effects in brain tumors and amplifications are associated with different tumor groups in brain tumors.(53) That is, recapitulation of *MYCN* expression in preclinical models could be vital depending on the specific research question or tumor grade of interest. The exclusive significant correlation of these transcription factors by preclinical model type underscores the importance in careful model selection.

To better understand the specific genes driving correlation (and anti-correlation) of models to their tissue and disease context, we found the genes with the highest correlation, lowest correlation, and anti-correlation between each GBM model and brain tumors and brain non-diseased tissues (Figure 5). Functional enrichment analyses by similar studies have previously focused on using differentially expressed genes in cell lines compared to tumors as input.(9,10,23) While this is important to highlight the biological terms that are enriched just in preclinical models versus just in patient tissues, we also wanted to identify gene programs that may be particularly well represented in a model, by looking at overrepresentation of highly correlated genes between GBM-derived models and brain tumor or non-diseased brain tissue. Interestingly, some of the top enriched terms for cell lines by positive correlation to non-diseased brain are related to primary cilia, oxidative phosphorylation, and cellular response to heat shock in brain tumors (Figure 5C). In PDXs, we observed top enriched terms for positively correlated genes for exercise-induced circadian rhythm for brain tumors, cell-cell signaling, and *ZNF148* transcription factor activity (Figure 5C). Genes with a negative correlation between cell lines and tumors were enriched for immune pathways, consistent with previous findings.(10,23) Interestingly, multiple metabolic pathways were negatively enriched between GBM PDXs and brain tumors as well as non-diseased brain tissues. While PDX models are preferred for drug testing because of their higher fidelity to patient tumor recapitulation and heterogeneity, they are lower throughput and more expensive than cell lines and our findings suggest they may not recapitulate all aspects of drug testing better than cell lines.

There are several limitations to this study. While we removed 66 cancer cell lines from our study that were known to be problematic in the Cellosaurus (i.e., evidence of contamination or misidentification), other cancer cell lines included may be misidentified or contaminated but not reported. Another limitation of our study is that while we assessed how genes associated with tumor purity and tissue-specific expression impact the global gene expression correlation between cancer cell lines and human tissues, future studies will need to assess if particular pathways or other gene sets are conserved or not and the impact of other tumor-specific characteristics like subtype or tumor grade. Finally, given the robust number of GBM cell lines and PDX gene expression profiles, our comparative analyses between cancer cell lines and PDXs were focused on these.

While we found that GBM PDX models better recapitulated tissue-specific gene expression than GBM cell lines when correlated with cancer and non-diseased brain tissues, this may or may not be true for cancer cell lines and PDX models derived from other tissues.

## Conclusion

Our findings 1) underscore the importance of using the full gene expression set measured when comparing preclinical models and tissues and 2) confirm that tissue-specific patterns are better preserved in GBM PDX models than in GBM cell lines. The latter is not surprising, as these cancer cell lines and PDX models are from GBM, so restricting tissue-specific gene expression of non-brain tissues should reduce that correlation. However, given the important role of preclinical models in oncology research, our study suggests that, at least for brain tumor research, careful consideration of what aspects of tumor gene expression are reproduced in cancer cell lines compared to PDXs may be critical. Additionally, our study is the first to our knowledge to compare gene expression correlation of preclinical models across matched tumor and non-diseased tissues. Future studies can build on these findings to determine the specific pathways and gene sets recapitulated by particular preclinical models to facilitate model selection for a given study design or goal.

## Supporting information

Supplementary Table 1: Problematic Samples Based on Metadata Mislabeling or Possible Contamination.

Supplementary Table 2: Tissue Types Common Between Data Sources.

Supplementary Table 3: Cancers Represented Between Data Sources.

Supplementary Table 4: Median Correlation of Each Individual Model Sample to Each Tissue Grouping.

Supplementary Table 5: Significant Pathway Terms.

## Acknowledgments

We would like to thank members of the Lasseigne Lab for their thoughtful feedback.

## Supplementary Figures

**Supplementary Figure 1.**
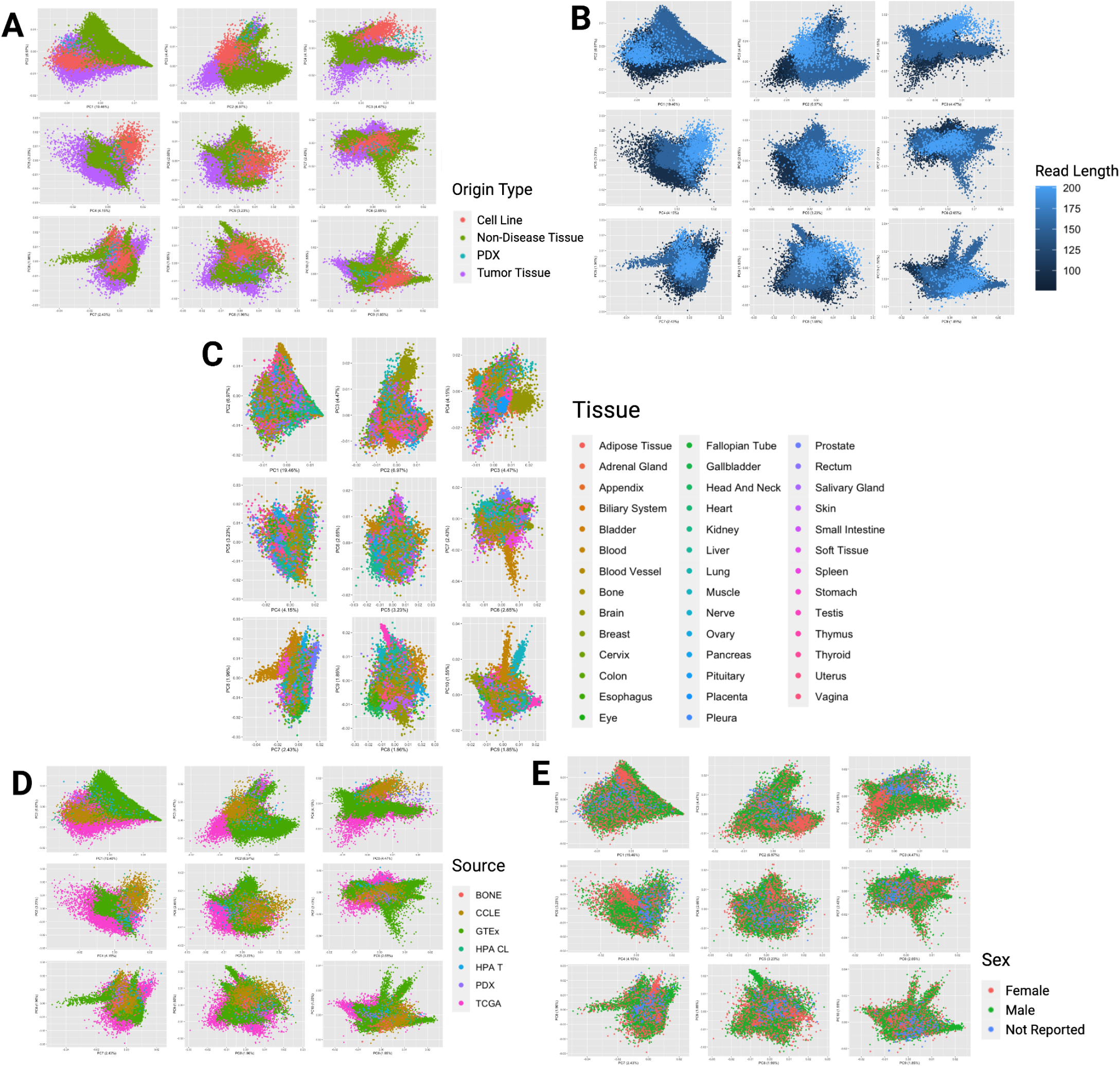
**A.** Principal component analysis scatterplot with density rug plot of gene expression profiles from cancer cell lines (red), GBM PDX models (teal), non-diseased tissue (green), and tumor tissue samples (purple) in principal components 1-10. **B**. Principal component analysis scatterplot with density rug plot of gene expression profiles from the same groups colored by read length. **C**. Principal component analysis scatterplot with density rug plot of gene expression profiles from the same groups colored by tissue type, regardless of disease state. **D**. Principal component analysis scatterplot with density rug plot of gene expression profiles from the same groups colored by the data resource samples originated from. **E**. Principal component analysis scatterplot with density rug plot of gene expression profiles of the same groups colored by sex of the individual that samples were originally derived from.

**Supplementary Figure 2.**
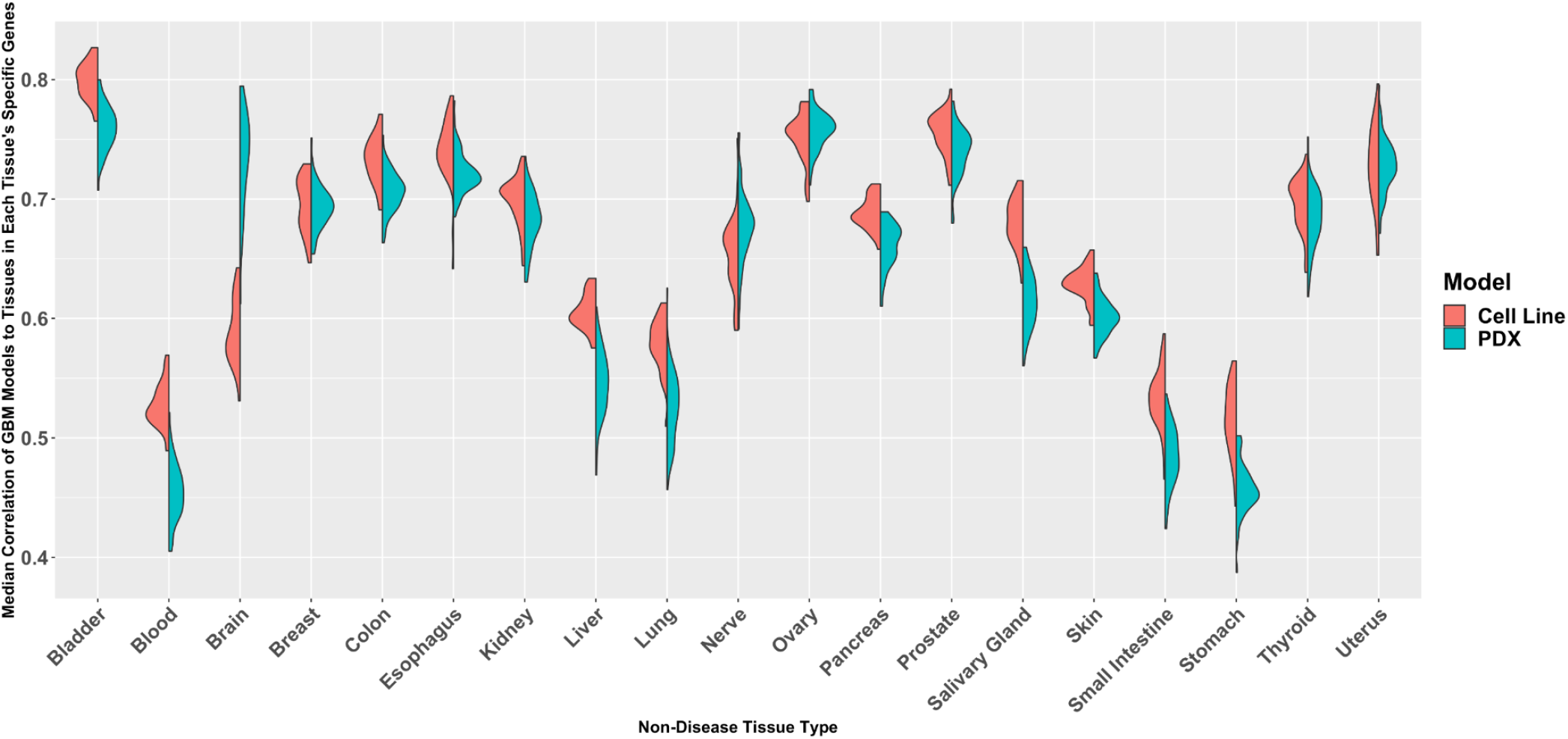
Split violin plots of the correlation of each GBM cancer cell line and PDX sample to each non-diseased tissue type by that tissue’s tissue-specific genes.

**Supplementary Figure 3.**
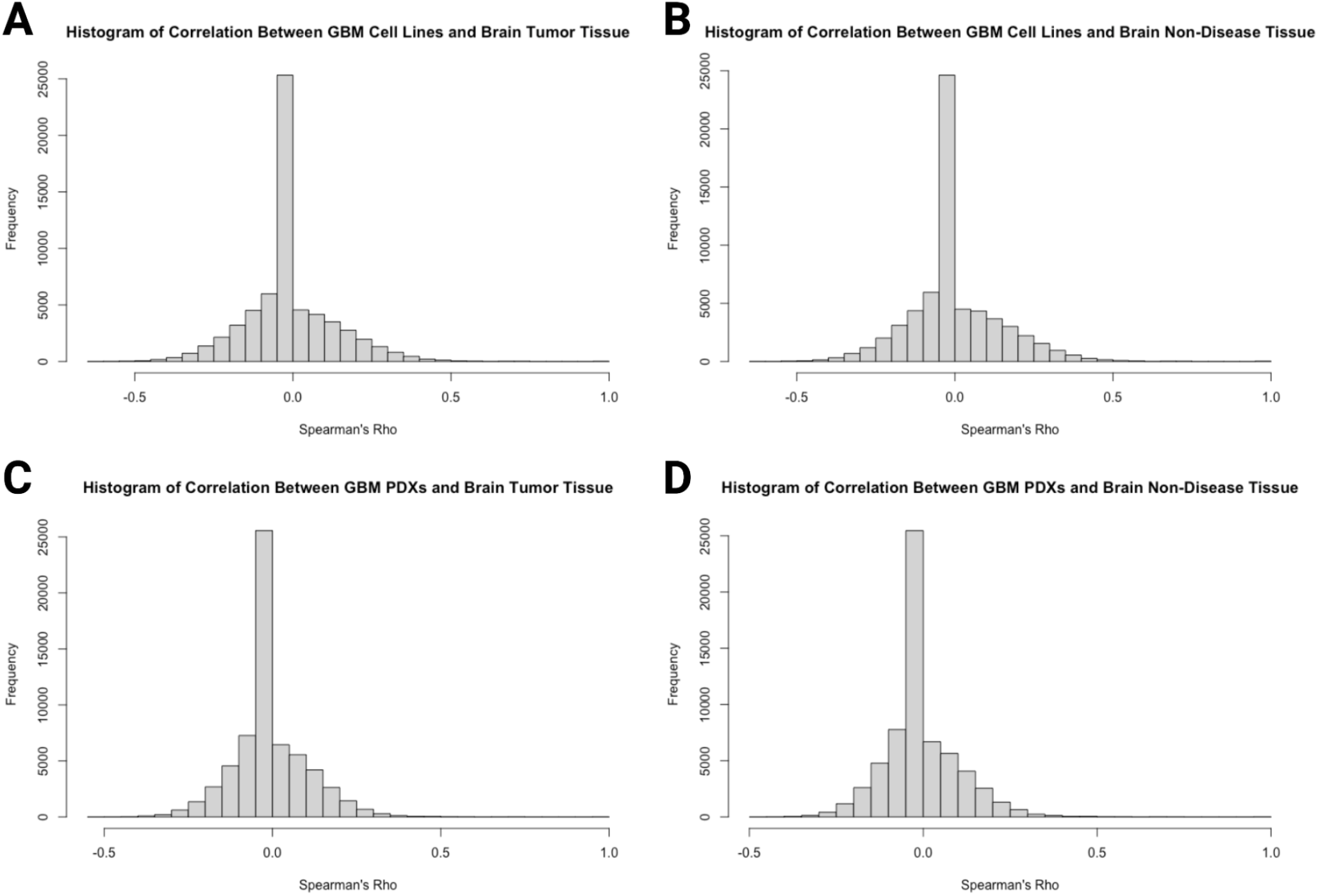
Histogram of the distribution of Spearman’s correlation rho between **A**. GBM cell lines and brain tumor tissue, **B**. GBM cell lines and brain non-diseased tissue, **C**. GBM PDXs and brain non-diseased tissue, and **D**. GBM PDXs and brain tumor tissue, respectively.

**Supplementary Figure 4.**
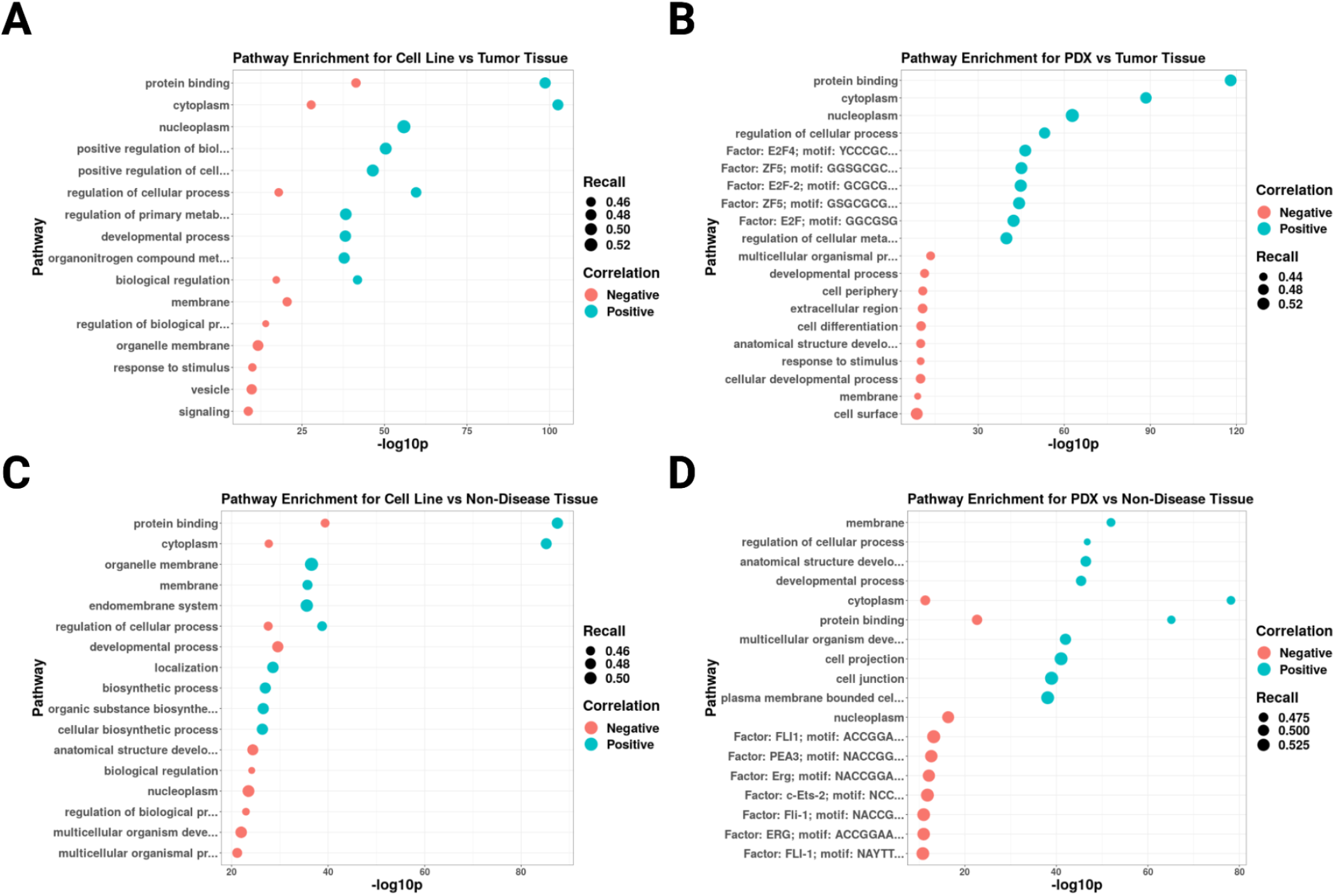
Dot plots of the most significant (top 10 if at least 10 were significant) pathways represented in insignificant genes with either positive (blue) or negative (red) correlation between **A**. GBM cell lines and brain tumor tissue, **B**. GBM PDXs and brain tumor tissue, **C**. GBM PDXs and brain non-diseased tissue, and **D**. GBM cell lines and brain non-diseased tissue, respectively. Dots are sized by recall or the proportion of genes from a specific term represented in the data divided by the total genes possible for that term.

### Supplementary Tables

[SupplementaryTable1_ProblematicSamples.xlsx]:

Supplementary Table 1: Problematic Samples Based on Metadata Mislabeling or Possible Contamination.

[SupplementaryTable2_CommonTissueTypes.xlsx]:

Supplementary Table 2: Tissue Types Common Between Data Sources. [SupplementaryTable3_CancersRepresented.xlsx]:

Supplementary Table 3: Cancers Represented Between Data Sources. [SupplementaryTable4_MedianCorrelations.xlsx]:

Supplementary Table 4: Median Correlation of Each Individual Model Sample to Each Tissue Grouping. [SupplementaryTable5_SignificantPathwayTerms.xlsx]:

Supplementary Table 5: Pathway Terms Represented in GOSt Results for Each Gene Set of Interest.

## Notes

**Funding Statement:** This work was supported in part by OD R03OD030604 (to BNL; supporting BNL, ASW, and JLF), R00HG009678 (to BNL; supporting BNL, EJW, AWS, JLF), an ACS-IRG (to BNL; supporting JLF), the UAB Lasseigne Lab Start-Up funds (to BNL; supporting BNL, ASW, EJW, and JLF), and the UAB Pilot Center for Precision Animal Modeling (C-PAM) 1U54OD030167 (supporting BNL and EJW). The funders had no role in the design of the study and collection, analysis, and interpretation of data and in writing the manuscript.

**Conflict of Interest Disclosure:** The authors have nothing to disclose.

https://zenodo.org/record/8044711

https://doi.org/10.5281/zenodo.7813907

https://hub.docker.com/r/lizzyr/rstudio_modelselection

### Competing Interest Statement

The authors have declared no competing interest.

### Summary of Updates

We have further clarified the use of varying gene set sizes and expanded the Discussion to better place our work in the larger context of the field.

